# Sudden heart rate reduction upon optogenetic release of acetylcholine from cardiac parasympathetic neurons in perfused hearts

**DOI:** 10.1101/451450

**Authors:** Angel Moreno, Kendal Endicott, Matthew Skancke, Mary Kate Dwyer, Jaclyn Brennan, Igor R Efimov, Gregory Trachiotis, David Mendelowitz, Matthew W Kay

**Affiliations:** Department of Biomedical Engineering, The George Washington University, 800 22nd Street NW, Suite 5000, Washington, DC 20052, USA; Department of Pharmacology and Physiology, The George Washington University, 2300 Eye Street NW, Suite 640, Washington, DC 20037, USA; Division of Cardiothoracic Surgery, Veterans Affairs Medical Center, 50 Irving St. NW., Washington, DC 20422, USA

**Keywords:** neurocardiac, cardiac optogenetics, autonomic nerve activation, cardiac parasympathetic tone, excised perfused hearts

## Abstract

The balance of sympathetic and parasympathetic tone provides exquisite control of heart rate and contractility and has also been shown to modulate coronary flow and inflammation. Understanding how autonomic balance is altered by cardiac disease is an active area of research and developing new ways to control this balance is providing insights into disease therapies. However, achieving acute neuron-specific stimulation of autonomic neurons can be difficult in experiments that measure the acute effects of nerve stimulation on the heart. Conventional electrical and pharmacological approaches can be spatially and temporally non-selective. Cell-specific expression of light-activated channels (channelrhodopsin, ChR2) is a powerful approach that enables control of the timing and distribution of cellular stimulation using light. We present such an optogenetic approach where parasympathetic cardiac neurons are selectively photoactivated at high temporal precision to initiate cholinergic-mediated slowing of heart rate.

Mice were crossbred to express ChR2 in peripheral cholinergic neurons using Cre-Lox recombination driven by a choline acetyltransferase (ChAT) promoter. Hearts from adult mice were excised, perfused, and the epicardium was illuminated (peak 460-465nm) to photoactivate ChR2. In one set of studies, hearts were illuminated using a large-field LED light source. In other studies, a micro LED was placed on the right atrium to selectively illuminate the junction of the superior vena cava and right atrium. The ECG was acquired before, during, and after tissue illumination to measure changes in heart rate. Upon illumination, hearts exhibited sudden and dramatic reductions in heart rate with restoration of normal heart rate after cessation of illumination. Delays in atrioventricular conduction were also observed. Heart rate reductions at the highest irradiance levels were similar to heart rate reductions caused by application of bethanechol (10µM) or acetylcholine (800µM). Atropine (50nM) completely blocked the effect of ChR2 photoactivation, confirming cholinergic mediation.

Optogenetic activation of intrinsic parasympathetic neurons reduced heart rate in an immediate, dose-dependent fashion, resembling the slowing of sinus rate in response to acetylcholine. Our results demonstrate a new approach for controlling parasympathetic modulation of cardiac function by selectively activating the endogenous release of acetylcholine from intrinsic cardiac cholinergic neurons.

**Key Message:** Optogenetic photoactivation of intrinsic cardiac neurons provides immediate, tissuespecific stimulation with minimal cross-reactivity. Our results demonstrate that selective expression of channelrhodopsin within cardiac cholinergic neurons enables photoactivated release of acetylcholine, thereby instantaneously slowing sinus rate and altering atrioventricular conduction. This provides for indepth examination of the endogenous interplay between cardiac autonomic neurons and the functional outcomes of downstream post-synaptic receptor activation.

## 1. Introduction

The balance of sympathetic and parasympathetic tone provides exquisite control of heart rate and contractility and has also been shown to modulate coronary flow (Kovach et al., 1995; Reid et al., 1985) and inflammation after myocardial injury (Calvillo et al., 2011; Garrott et al., 2017). Understanding how autonomic balance is altered by cardiac disease is an active area of research and developing new ways to control this balance is providing insights into disease therapies. In the laboratory, autonomic tone has been modulated by pharmacological activation of cardiomyocyte membrane receptors and by electrical stimulation of sympathetic ganglia or the vagus nerve. These methods are effective and initiate profound changes in heart rate (DeWitt et al., 2016; Ng et al., 2001). However, measuring changes in cardiac function that result from the release of neurotransmitter from a specific population of cardiac neurons (anatomic-functional probing) can be difficult because of the difficulty in achieving neuron-specific stimulation.

Optogenetics enables spatially and temporally specific stimulation of distinct cell types using light (Entcheva, 2013). Photostimulation of neurons expressing the light-gated cation channel channelrhodopsin (ChR2) initiates the immediate release of endogenous neurotransmitters (Deisseroth et al., 2006). Previous studies in brain slices having cholinergic neurons that express ChR2 demonstrated temporal control in the photoactivated release of acetylcholine (ACh) (Hedrick et al., 2016; Nagode et al., 2011; Ren et al., 2011). For the heart, cell-specific expression of ChR2 eliminates the need for complex anatomic dissections for electrical stimulation or the administration of circulating agonists that ubiquitously activate membrane receptors. With optogenetics, either branch of the cardiac autonomic nervous system could be selectively stimulated, with relative ease, in a temporally and regionally specific manner. For example, photoactivation of sympathetic neurons within the myocardium of perfused hearts was accomplished by illuminating the epicardium with blue light (Wengrowski et al., 2015). In those studies, the expression of ChR2 was targeted to catecholaminergic neurons using a Cre-Lox approach where the expression of Cre recombinase was promoted by tyrosine hydroxylase (TH), an enzyme involved in the production of norepinephrine.

The goal of the present studies was to demonstrate optogenetic photoactivation of intrinsic cardiac parasympathetic neurons and to measure the resulting changes in cardiac electrical activity. Our hypothesis was that hearts having ChR2 selectively expressed within cells that contain choline acetyltransferase (ChAT) could be photoactivated to induce changes in cardiac function consistent with muscarinic (M2) receptor activation. This is because ACh is produced by ChAT through the combination of acetyl-CoA with a free choline (Wessler et al., 1999) and is stored in synaptic vesicles within axon varicosities. Upon axonal depolarization, and the resulting increase in cytosolic calcium concentration, the vesicles dock with the membrane to release ACh, which binds to cardiomyocyte M2 receptors. This activates inhibitory G proteins (Irisawa et al., 1993) to modulate cardiomyocyte membrane currents that hyperpolarize the cell, slow spontaneous membrane depolarization of the sinoatrial node (SAN), and increase atrioventricular (AV) conduction time (Imaizumi et al., 1990). ACh also activates G protein-gated inwardly rectifying K^+^ channels (GIRK, also known as I_KACh_), which are thought to be a pivotal contributor to parasympathetic regulation of heart rate (Lee et al., 2018).

Previous studies have shown that electrical stimulation of the right and left vagus nerves induce immediate reductions in sinus rate and increases AV conduction time (Levy and Zieske, 1969; Ng et al., 2001). This is consistent with the release of ACh from parasympathetic cholinergic varicosities within those nodal regions (Imaizumi et al., 1990; Roy et al., 2014). Furthermore, local electrical stimulation of specific cardiac vagal ganglia in the fat pads, or their selective ablation, has revealed that the SAN and the AV node are innervated by distinct cholinergic axons (Randall et al., 1986; Sampaio et al., 2003). In rats, stimulation of the ganglion at the junction of the right superior vena cava and right atrium slowed heart rate with little effect on AV conduction while stimulation of the ganglion at the junction of the inferior pulmonary veins and left atrium slowed AV conduction without eliciting bradycardia (Sampaio et al., 2003). That study is an elegant example of anatomic-functional probing of the intrinsic cardiac nervous system to reveal new insights that could not be obtained by either administering exogenous neurotransmitters or by direct stimulation of the vagus nerve.

The studies presented here demonstrate an optogenetic approach for anatomic-functional probing of the parasympathetic nervous system that can be used with conventional excised perfused heart preparations. This is a corollary approach to our previous work detailing photoactivation of sympathetic axons (Wengrowski et al., 2015). Our results also demonstrate that cholinergic neurons remain active and viable following the severance of pre-ganglionic axons and, importantly, that the parasympathetic pathway, from post-ganglionic neuron stimulation to M2 receptor activation, can be successfully controlled using light in excised perfused hearts.

## 2. Methods

### 2.1 Mice with parasympathetic neurons that express ChR2

Approval for all animal protocols was obtained from the George Washington University’s Animal Care and Use Committee and followed the National Institute of Health’s Guide for the Case and Use of Laboratory Animals. Transgenic mice were crossbred to express ChR2 in cardiac parasympathetic neurons using Cre-Lox recombination and a ChAT promoter. One parent (Jackson Labs stock #006410) had homozygous expression of a ChAT promoter to direct the expression of Cre recombinase (Rossi et al., 2011) while the other parent (Jackson Labs stock #012569) had homozygous ChR2&EYFP fusion protein expression dependent upon Cre expression (Madisen et al., 2012). The enhanced yellow fluorescent protein (EYFP) enabled ChR2 expression to be visualized without fluorescent antibody staining. Genotyping of tail snips (Transnetyx) confirmed the genotype of the offspring (ChAT-Cre-ChR2&EYFP).

### 2.2 Whole-mount preparations and confocal microscopy

Multiphoton confocal imaging of excised right atria confirmed that ChR2&EYFP expression was localized to cells containing ChAT. In those studies, mice were anesthetized, cervically dislocated, the heart was rapidly excised, the aorta was cannulated, and the heart was flushed with phosphate buffered saline (PBS) containing heparin. Using a dissecting scope, the base of the heart, including the atria, was separated from the ventricles along the atrial-ventricular ridge and attached to a thin piece of polydimethylsiloxane (PDMS, Sylgard 184) (Rysevaite et al., 2011). The sample was then fixed with 4% paraformaldehyde in PBS for 2 hours. The heart was then thoroughly washed with PBS (3×2min, 2×15min, 1×30min) and blocked for two hours with 2% bovine serum albumin (BSA) (Sigma-Aldrich), 1% Triton X-100 (Sigma-Aldrich) in PBS. Next, the sample was placed in goat anti-ChAT primary antibody (EMD Millipore AB144P; 1:100 with 2% BSA) overnight at 4°C, with agitation. The sample was washed to remove all of the unbound primary antibody (3×2min, 2×15min, 1×1hr, 1xovernight). Following this, the sample was incubated with an Alexa Fluor 647 conjugated donkey anti-goat secondary antibody (Jackson Immunoresearch 705-605-147; 1:300 with 2% BSA) overnight at 4°C, while shaking. The heart was then washed in PBS (3×2min, 2×15min, 1×1hr, 1xovernight) and imaged using a Leica TCS SP8 MP multiphoton confocal microscope with a dry 10X lens. The sample was illuminated with light at 514nm and 651nm to visualize EYFP and the Alexa Fluor antibody, respectively.

To further examine the expression of ChR2&EYFP in cardiac neurons, hearts were excised from ChAT-Cre-ChR2&EYFP mice (n=4) and the right atria (RA) were removed and placed in PBS at room temperature. Submerged atria were then positioned on the stage of the Leica multiphoton confocal microscope and imaged using a water emersion lens (25X 1.0 µm critical aperture). Samples were illuminated at 514 nm to excite EYFP. Z stack images were acquired from the atriocaval junction (AC), at the intersection of the RA and the superior vena cava (SVC), to visualize EYFP fluorescence in three dimensions.

### 2.3 Excised perfused heart preparations

Mice were anesthetized with ∼4% isoflurane. After confirming a surgical plane of anesthesia, hearts were rapidly excised, and the aorta was cannulated. Hearts were then retrogradely perfused via Langendorff at constant pressure between 60 and 80 mmHg using a modified Krebs-Henseleit solution (118 mM NaCl, 4.7 mM KCl, 1.25mM CaCl_2_, 0.57mM MgSO_4_, 25mM NaHCO_3_, 1.17mM KH_2_PO_4_, 6mM glucose). Perfusate was buffered to a pH between 7.35 and 7.45, warmed to 37°C, and oxygenated with 95% O_2_/5% CO_2_. Before beginning each protocol, perfused hearts were allowed to stabilize at normal sinus rhythm for at least 5 min. Hearts were then positioned with the RA/SVC junction face up (posterior side) for photoactivation. Table 1 lists the number of animals used in each perfused heart study.

**Table 1.** Number of animals used in each set of experiments. With the exception of wild-type mice, all animals expressed ChR2&EYFP in ChAT neurons, as verified by functional responses and/or whole-mount confocal fluorescence imaging.

**Number of animals used in each study**
| Study | n |
| --- | --- |
| **Spotlight LED (all intensities)** | **10** |
| *Bethanechol* | 4 |
| *Atropine* | 3 |
| **Micro LED** | **10** |
| *Atropine* | 6 |
| *Optical mapping* | 3 |
| **Photoactivation with intact Fat pad** | **3** |
| *Green light stimulation verification* | 3 |
| **Photoactivation desensitization** | **3** |
| **WT photoactivation test** | **5** |
| *Bethanechol* | 5 |
| **WT ACh** | **9** |

### 2.4 ECG measurements

Three needle electrodes were positioned in an Einthoven configuration around hearts to record the bath-conducted ECG (Figure 1A). The signal was amplified and filtered (2122i Bioamplifier, UFI) and acquired using a PowerLab unit (PowerLab 8/35, AD Instruments). The ECG and heart rate (RR interval) were monitored in real time (LabChart, AD Instruments). The time of the P-wave and PQ intervals were measured before, during, and after specific illumination protocols.

**Figure 1.**
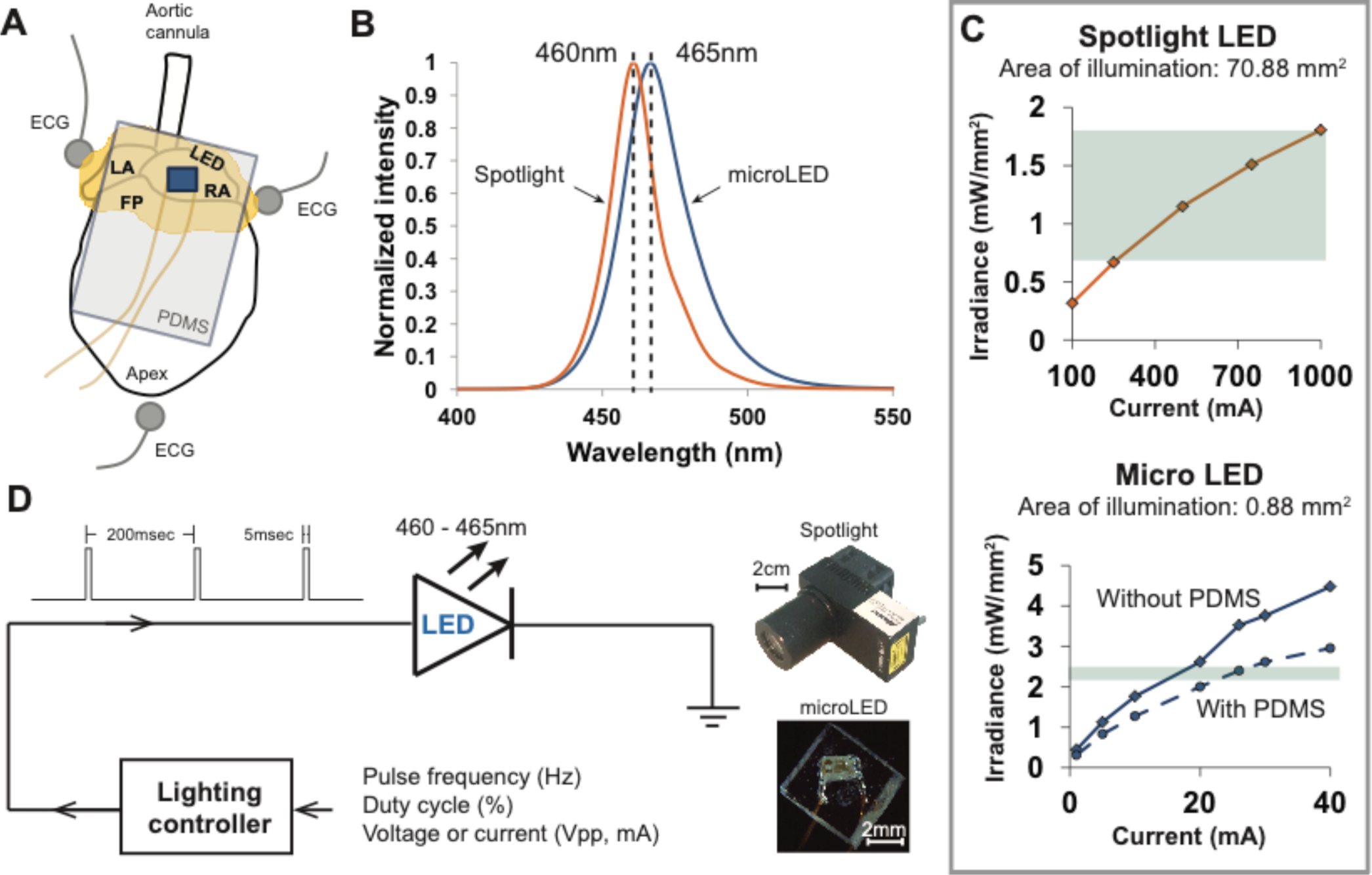
*Methods used to photoactivate ChR2 expressed within the cholinergic neurons of excised perfused mouse hearts* **A**: Arrangement of the heart (fat pads shown in yellow) during an experiment when photoactivating ChR2 using a PDMS-encapsulated micro LED, which is shown lying on top of the RA. Three electrodes recorded the bath-conducted ECG. **B:** The spectral distribution of light from the spotlight LED was centered at 460nm. The spectral distribution of light from the micro LED was slightly wider and centered at 465nm. **C**: The relationship between emitted light (irradiance) and LED source current for the two illumination devices used in the experiments. High source currents (250-1000mA) were required to generate sufficient irradiance (0.7-1.8mW/mm^2^) to activate ChR2 with the spotlight LED (top). The micro LED (bottom) provided ample irradiance (2.4mW/mm^2^) to activate ChR2 at low source currents (18-22mA). The range of irradiance used in the experiments is denoted by the green regions. **D**: Scheme for controlling the delivery of light from either light source. A lighting controller was programmed to deliver current to each LED as a pulse train at a specific frequency and duty cycle. The relative sizes of each LED source are shown on the right.

### 2.5 Photoactivation using a spotlight LED

Light from a high-power blue LED spotlight (Figure 1, 460 nm peak wavelength, Mightex Systems) was directed onto the right atrium and pulsed for 20 seconds (5 ms pulse width at 5 Hz). The current powering this spotlight could be varied to provide a range of optical irradiances, enabling the measurement of “dose-response” changes in heart rate (HR). Irradiances between 0.68 to 1.81 mW/mm^2^ (Figure 1C) were studied. This range was selected using results of previous studies (Wengrowski et al., 2015). After each illumination, light was withdrawn for 20 seconds and the heart was allowed to stabilize before applying the next illumination. The bath-conducted ECG was continuously recorded throughout each photoactivation protocol.

### 2.6 Photoactivation using a micro LED

Small solid-state LEDs (Figure 1) were used to focus excitation light within specific regions of the heart while also demonstrating that small LEDs could be effective in activating cardiac neurons that express ChR2. We chose surface-mount LEDs (Dialight 598 series, Vf 3.2V), with a peak wavelength at 465nm and maximum irradiance of 2.4mW/mm^2^ (Figure 1B&C), to illuminate the RA/SVC junction (Figure 1A). Given the small size (1.6mm x 0.8mm x 0.7mm) of these LEDs, significant illumination of adjacent areas of the heart was avoided, thereby eliciting only the response from targeted locations. To avoid electrical short circuits at the interface of the LED due to interaction with the surrounding electrolyte solution, the LEDs were encapsulated with PDMS (Sylgard 184). Due to the high resistivity of PDMS (2.9×10^14^ Ω.cm), this provided suitable thermal and electrical insulation. A thin layer of PDMS (15µm) was first applied to the front of the LED, where the light is emitted, to minimize light attenuation. A second thicker layer of PDMS was then applied to the rest of the LED chip (Figure 1D). Current was delivered to micro LED devices in a manner similar to that used for spotlight illumination.

### 2.7 Administration of muscarinic agonists and antagonists

The muscarinic agonists bethanechol (carbamyl-β-methylcholine chloride, Sigma) (10µM) and ACh (acetylcholine chloride, Sigma) (800µM) were administered to the perfusate as positive controls to measure the effects of ubiquitous activation of muscarinic receptors. This allowed the effects of photoactivation of cholinergic neurons to be compared to that of muscarinic activation by circulating agonists. To demonstrate cholinergic specificity of photoactivation, atropine (atropine sulfate, Sigma) (50nM), a muscarinic antagonist, was administered to the perfusate. Cessation of changes due to photoactivation were then measured using the spotlight and micro LED at irradiances of 1.81 and 2.4 mW/mm^2^, respectively; which provided the largest drop in HR without atropine.

### 2.8 Optical mapping experiments

To study sinus-mediated beats and atrio-ventricular conduction time, optical mapping of a voltage sensitive dye (di-4-ANEPPS) was performed before and after photoactivation using the micro LED. The dye (30 µL of 2.5mM stock diluted in 1 mL of perfusate) was injected into the aorta. Upon verification of sufficient staining, blebbistatin (circulating concentration of 5µM) was titrated into the perfusate to mechanically arrest hearts to prevent motion contamination of the optical signals. A high-power green LED light source with peak wavelength of 520 nm (UHP-Mic-LED-520, Prizmatix) illuminated the epicardial surface to energize the dye. Emitted light was long pass filtered (680nm) and imaged with a MiCAM Ultima-L CMOS camera (SciMedia, Costa Mesa, CA) with high spatial (100×100 pixels, 230±20 μm per pixel) and temporal (1,000-3,000 frames/sec) resolution. The resulting optical action potential (OAP) datasets were stored for subsequent analysis.

### 2.9 Data analysis and statistics

Data are presented as mean ± standard error of the mean. Statistical analyses were performed using analysis of variance, paired t-test and Bonferroni post-hoc testing using SPSS v24 with p<0.05 indicating significance. Optical mapping datasets were pre-processed using custom MATLAB algorithms to increase contrast, normalize signals, and identify times of OAP depolarization.

## 3. Results

### 3.1 Tissue whole-mount fluorescence microscopy

Neurons within right atria that expressed ChAT were labeled via immunohistochemistry with Alexa Fluor 647 to provide visual confirmation that ChAT and ChR2&EYFP expression were localized within the same axons (Figure 2A). Fluorescence overlap confirmed selective expression of ChR2&EYFP within cholinergic neurons. Right atrial nerve bundles that expressed ChR2&EYFP were also imaged at high resolution within the region of the atriocaval (AC) junction (Figure 2B). An extensive intertwining network of nerve fibers was observed, indicating robust expression of ChR2&EYFP in this animal model. Nerve bundles contained axons with diameters between 3 and 5.4 µm.

**Figure 2.**
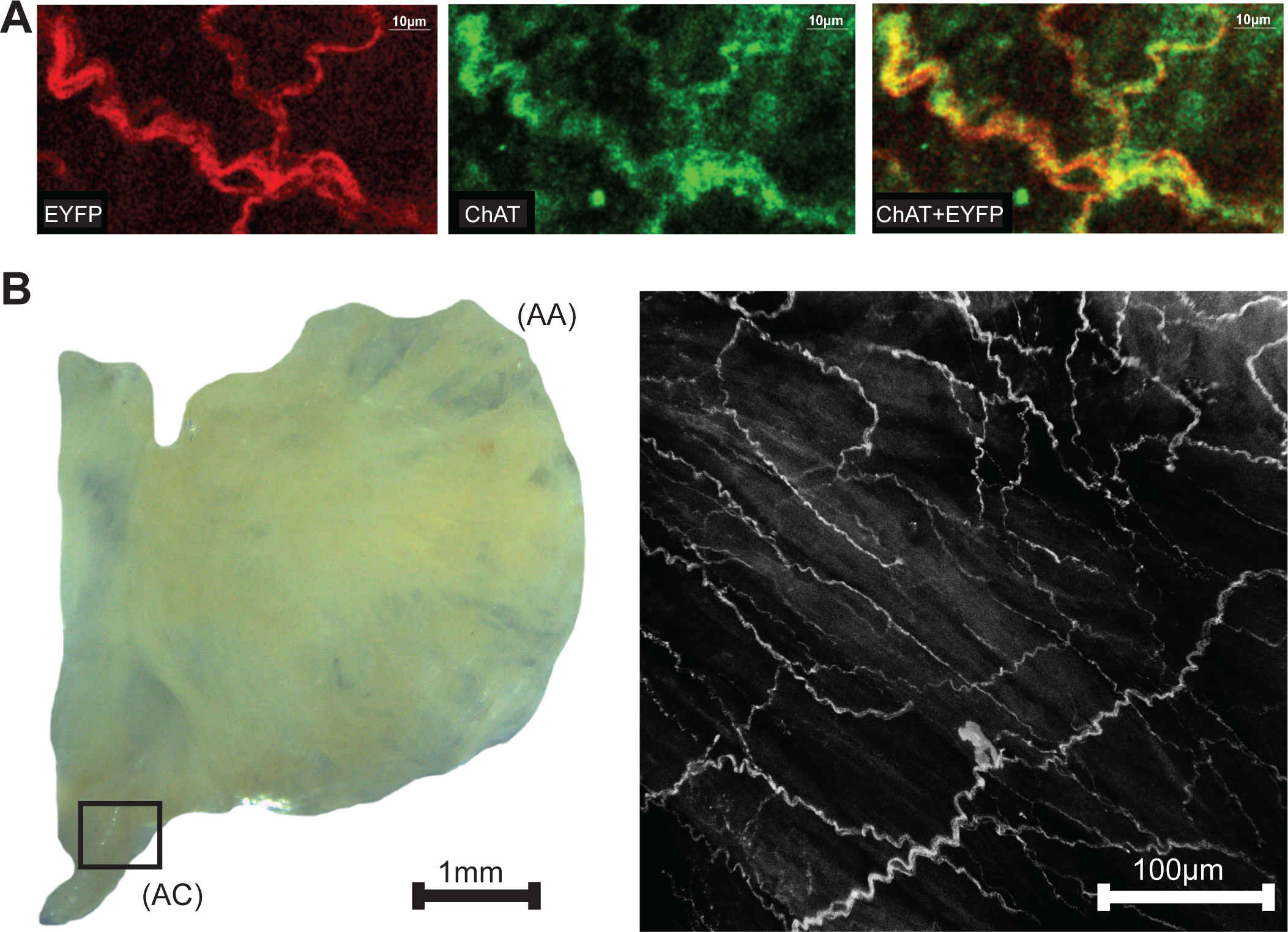
*Images showing colocalization of ChAT with ChR2&EYFP within the nerve bundles of the right atrium.* **A**: The fluorescence of EYFP (left, 514nm excitation) and Alexa Fluor (middle, 651nm excitation) were imaged within the RA. Fluorescence overlap (right) confirmed selective expression of ChR2&EYFP within cholinergic neurons. **B:** Bright field image (left) of a typical excised right atrium used for multiphoton fluorescence microscopy of neurons expressing ChR2&EYFP. AC: atriocaval junction; AA: atrial appendage. Extensive intertwining networks of nerve fibers expressing ChR2&EYFP are shown on the right for a region of the AC junction, denoted as the box shown in the bright field image.

### 3.2 Heart rate reduction after photoactivation

Following cannulation, perfused hearts (n=10) stabilized at 320 ± 17 bpm. Immediately prior to illumination, average baseline HRs for each irradiance were 317 ± 18 bpm for 0.68 mW/mm^2^, 321 ± 16 bpm for 1.15 mW/mm^2^, and 319 ± 17 bpm for 1.81 mW/mm^2^ (Mean ± SE). During illumination, HR demonstrated an immediate, dose-dependent reduction. Typical ECG signals before and during illumination are shown in Figure 3A. Average HRs during each irradiance level were (Figure 3B): 240 ± 22 bpm for 0.68mW/mm^2^, 208 ± 15 bpm for 1.15mW/mm^2^, and 175 ± 14 bpm for 1.81 mW/mm^2^. Upon cessation of 20 seconds of illumination, average HRs returned to baseline after 5-10 sec (315 ± 20, 318 ± 16, and 326 ± 19 bpm for 0.68mW/mm^2^, 1.15mW/mm^2^, and 1.81 mW/mm^2^, respectively) regardless of the irradiance. Long-term photoactivation studies were performed where hearts (n=3) were continuously illuminated for specific time intervals (5, 10, 30 and 60 minutes). Drops in HR were maintained for at least 30 minutes and returned to baseline after cessation of illumination (Figure 3C, top). During 60 minutes of continuous illumination, desensitization was observed after 35 minutes where HR started to rise and at 10 minutes after illumination HR remained low (Figure 3C, bottom).

**Figure 3.**
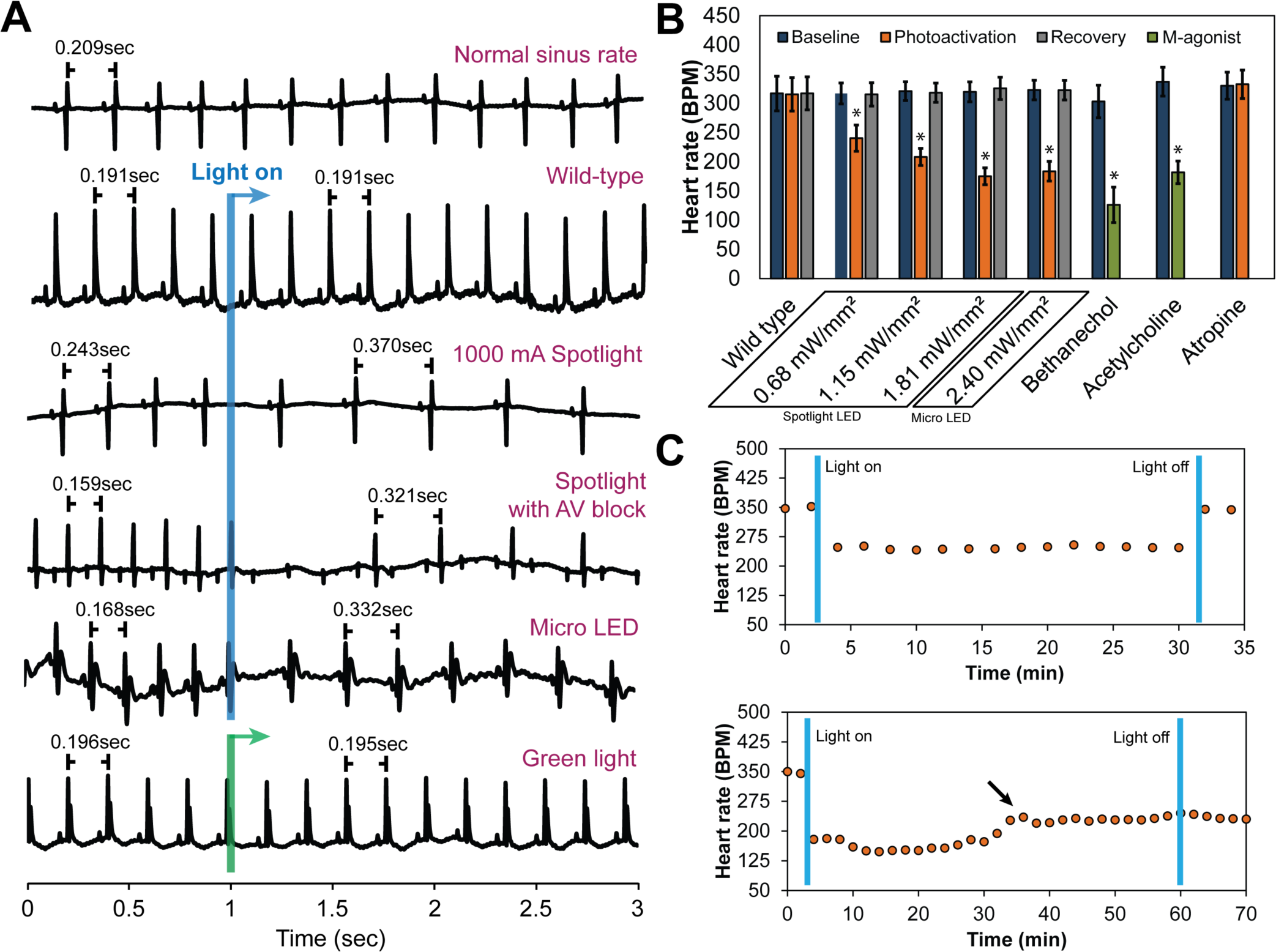
*The effect of cholinergic neuron photoactivation on sinus rate and AV conduction in perfused hearts from ChAT-Cre-ChR2&EYFP mice.* **A**: Six representative ECG signals demonstrate the effect of cholinergic neuron photoactivation on R-R interval and AV conduction. From top to bottom: 1) Typical baseline ECG before photoactivation. 2) Typical ECG at the onset of illuminating wild-type mouse hearts with the spotlight LED. No change in the ECG was observed. 3) Typical ECG at the onset of photoactivation using the spotlight LED at the highest source current. RR interval increased immediately. 4) Representative ECG showing AV block when illuminating the RA and AV node with the spotlight LED at the highest source current. Second-degree AV block was observed. 5) Typical ECG at the onset of photoactivation using the micro LED placed on top of the RA. RR interval increased immediately. 6) Representative ECG showing no change in HR at the onset of green light (peak 520nm) illumination of the RA of ChAT-Cre-ChR2&EYFP mouse hearts. **B**: Average HR changes (n=10 hearts) at baseline, during photoactivation of the RA at four different irradiances (with either the spotlight or micro LED), and after cessation of illumination. Average HR changes are also shown for wild-type mice with no ChR2 expression (n=5 hearts) and for the muscarinic agonists bethanechol (10µM, n=9 hearts) and acetylcholine (800µM, n=9 hearts). The muscarinic antagonist atropine (50nM, n=9 hearts) completely blocked changes in HR during photoactivation by the spotlight (1.81 mW/mm^2^) and the micro LED (2.4 mW/mm^2^). **C:** Robustness and stability tests were performed to assess the effect of prolonged photoactivation (n=3 hearts). **Top**: Change in HR during 30 min of photoactivation. HR returned to baseline upon cessation of illumination. **Bottom**: Change in HR during 60 min of photoactivation. The arrow indicates that the potency of photoactivation was reduced after 35-40 min of illumination. HR did not return to baseline until 20 min after cessation of illumination (data not shown).

As expected, wild-type animals with no ChR2 expression (n=5) demonstrated no response to LED illumination, regardless of the irradiance (Figure 3A). This confirmed that changes in HR were not caused by the direct effects of illumination, such as heating. In fact, changes in epicardial temperature during illumination were <0.03°C.

In a subset of studies (n=5), spotlight illumination of the RA near the AV node slowed HR and also induced AV block (Figure 3A). This occurred in both a 2:1 and 3:1 ratio for atrial to ventricular conduction, respectively. A prolonged PQ interval was observed during photoactivation that induced AV block (56.6 ± 3.6 vs 59.7 ± 3.8 ms, p=0.004) but no difference in the duration of atrial depolarization (P wave duration, 12.7 ± 1.3 vs 13 ± 1.3 ms, p=0.356) was evident. At cessation of illumination, AV block was relieved, and hearts immediately returned to normal sinus rhythm.

Immediate HR reductions (n=10 hearts) were also observed using the micro LED (2.4 mW/mm^2^) (Figure 3). HR was 323 ± 17 bpm immediately prior to illumination with the micro LED. During illumination, average HR dropped to 184 ± 17 (Figure 3B). Average HR after cessation of illumination was 322 ± 17 bpm. AV block was not observed when the micro LED was directed toward the SA node (Figure 3A). In the absence of AV block, no differences in atrial depolarization (P wave) duration and PQ interval were observed before or during illumination (16 ± 2.4 vs 17 ± 2.3 ms, p=0.117 and 54 ± 4.5 vs 53 ± 3.9 ms, p=0.298, respectively). In a separate group (n=3 hearts), the epicardial fat pads were carefully preserved during heart excision and cannulation. The average results of photoactivation in that subset of hearts was the same as that of all other hearts.

Bonferroni post-hoc testing of each irradiance indicated that all levels had statistically significant HR reductions (Figure 3B) compared to baseline HR before photoactivation (Table 2). Furthermore, irradiances of 1.15 (35%, p<0.001), 1.81 (45%, p<0.001) and 2.4 mW/mm^2^ (43%, p<0.001) all had a larger impact on HR compared to 0.68 mW/mm^2^ (24%, p<0.001). There was no statistically significant intergroup difference in HR reductions between irradiances of 1.15, 1.81 and 2.4 mW/mm^2^.

**Table 2.**
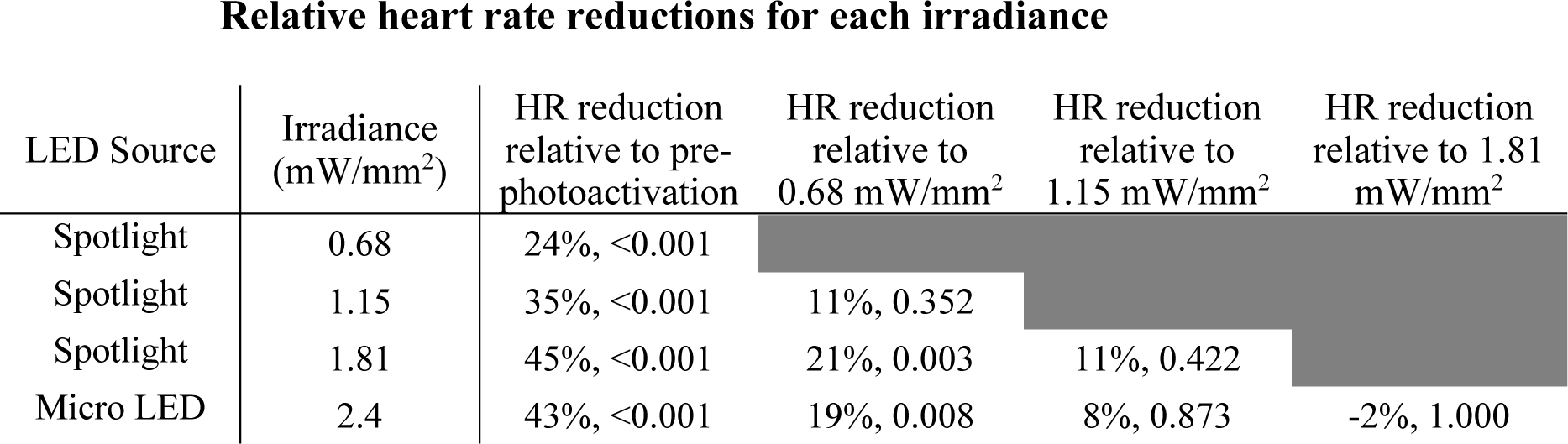
Percent HR reductions following cholinergic neuron photoactivation at each irradiance were analyzed using Bonferroni post-hoc comparisons. The third column shows the average percentage that HR was reduced during photoactivation relative to pre-photoactivation. Columns 4-6 compare HR reductions during photoactivation at each irradiance. Note the statistically significant difference in reduction for all irradiance levels compared to pre-photoactivation but no statistically significant difference between HR reductions at irradiance levels of 1.15, 1.81 and 2.4 mW/mm^2^.

| LED Source | Irradiance (mW/mm<sup>2</sup>) | HR reduction relative to pre-photoactivation | HR reduction relative to 0.68 mW/mm<sup>2</sup> | HR reduction relative to 1.15 mW/mm<sup>2</sup> | HR reduction relative to 1.81 mW/mm<sup>2</sup> |
| --- | --- | --- | --- | --- | --- |
| Spotlight | 0.68 | 24%, <0.001 |  |  |  |
| Spotlight | 1.15 | 35%, <0.001 | 11%, 0.352 |  |  |
| Spotlight | 1.81 | 45%, <0.001 | 21%, 0.003 | 11%, 0.422 |  |
| Micro LED | 2.4 | 43%, <0.001 | 19%, 0.008 | 8%, 0.873 | -2%, 1.000 |

### 3.3 Effects of muscarinic agonists and an antagonist

Bethanechol (10µM) produced a long-term transient reduction in HR from 302 ± 27 to 130 ± 10 bpm (Figure 3B) (n=9 hearts). Unpaired t-tests revealed no significant difference in HR reduction following bethanechol administration (p=0.09) between hearts of either wild type mice or ChAT-Cre-ChR2&EYFP mice. Bethanechol prolonged PQ interval (51 ± 4 vs 57 ± 5 ms, p<0.001) but did not modify the p-wave duration (12.1 ± 1.1 vs 12.5 ± 1.5 ms, p=0.576). ACh (800µM) was administered to the hearts of wild type mice (n=9), resulting in a long-term transient HR reduction from 337 ± 25 bpm to 182 ± 19 bpm (Figure 3B), significant AV delays (40.4 ± 1.6 vs 45.3 ± 2.8 ms, p<0.001), and preservation of p-wave duration (13.1 ± 0.7 vs 12.8 ± 0.9 ms, p=0.32). The observation that ACh did not alter the time of atrial depolarization is consistent with previous findings that ACh did not alter conduction velocity in canine right atria (Schuessler et al., 1990).

Atropine (50nM) was administered to the hearts of ChAT-Cre-ChR2&EYFP mice (n=9) and completely blocked the effects of photoactivation (330 ± 23 before illumination vs. 332 ± 24 bpm after illumination) (Figure 3B). No AV delay or atrial depolarization differences were observed before photoactivation with atropine or during photoactivation with atropine (54.8 ± 2.6 vs 51.6 ± 3.3 ms, p=0.055 and 12.8 ± 0.9 vs 11.9 ± 0.8 ms, p=0.134, respectively).

### 3.4 Optical mapping

Hearts were optically mapped (n=3) before and after photoactivation to study sinus-mediated beats and to measure the delay between the early site of atrial depolarization and the early site of LV depolarization (Figure 4). The earliest site of activation that was mapped within the atria and LV was used to compute this AV delay. Micro LED illumination lengthened AV delay by 6 ms but did not appear to alter the site of early LV depolarization. Bethanechol lengthened AV delay by 8 ms but did not appear to alter the site of early LV depolarization. Although there is minor spectral overlap in the light from the green 520 nm spotlight used for optical mapping and the absorption spectrum of ChR2 (Nagel et al., 2003), there was no measurable response that resulted from the green light. This was verified during control studies (n=3) where no change in HR was observed for the hearts of ChAT-Cre-ChR2&EYFP mice when the 520 nm spotlight used for optical mapping was used to illuminate the RA at the same duty cycle and frequency as that used for the 460 nm spotlight and the micro LED (Figure 3A).

**Figure 4.**
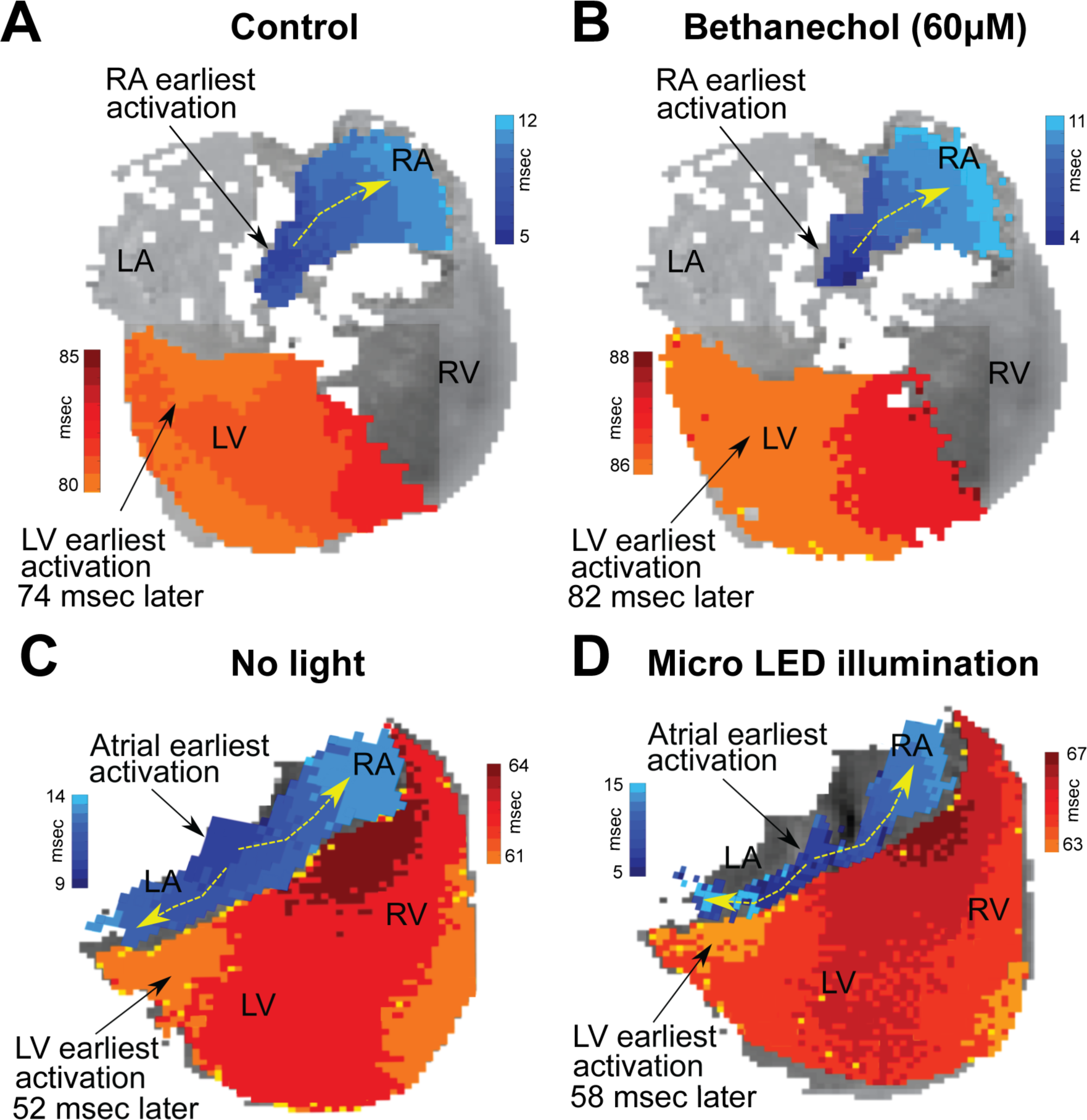
*Changes in conduction time between the atria and the ventricles induced by either bethanechol or photoactivation.* Hearts (n=3) were optically mapped to observe action potential propagation and to measure delays in conduction time between the atria and the ventricles, which was computed as the time difference between the earliest site of activation in the left ventricle (LV) and the earliest site of activation within the right atrium (RA). The PDMS-encapsulated micro LED was positioned on the base of the heart and directed toward the AC junction. **A**: RA and LV early sites of activation mapped before administering bethanechol (60µM). **B**: Bethanechol lengthened AV delay by 8 msec but did not appear to alter the site of early LV depolarization. **C**: RA and LV early sites of activation mapped before photoactivation. **D**: Photoactivation using micro LED illumination lengthened AV delay by 6 ms but did not appear to alter the site of early LV depolarization.

## 4. Discussion

Insights into cardiac autonomic neuroanatomy have been provided by early reports from Randall et al. (Randall et al., 1986) and subsequently improved in studies by Sampaio et al.(Sampaio et al., 2003) and Pauza et al.(Rysevaite et al., 2011). These groups successfully localized cardiac vagal ganglia and traced the parasympathetic neurons synapsing within the SA and AV nodes (Pauza et al., 2013; Sampaio et al., 2003). Observations from those studies indicate that the SA node is innervated by parasympathetic fibers originating from ganglia on both sides of the heart, but with a higher concentration of axons originating from right sided ganglia. Our work presents a novel approach to optogenetically drive endogenous parasympathetic release of ACh from those neurons to elicit downstream changes in cardiac function through the preservation of postganglionic parasympathetic neurons in excised hearts, either with or without intact fat pads. The specificity of cholinergic neuron photoactivation was confirmed with muscarinic receptor agonists and antagonists and verified with optical mapping demonstrating a lengthening of conduction time between the atria and ventricles. Photoactivation maintained low heart rates during short stimulation periods and for up to 30 min of continuous illumination; however, gradual decline of parasympathetic response during stimulation was observed after approximately 35 min. This was possibly a result of desensitization of the postsynaptic muscarinic receptors, progressive decrease in the release of ACh, or a fading of effector response (Martin et al., 1982; S. et al., 1993).

The irradiance-dependent response that we observed supports and extends previous results of reduced heart rates following electrical stimulation of the vagus nerve in animal models (Brack et al., 2004; Choate and Feldman, 2003; Degeest et al., 1965; Levy and Zieske, 1969). Heart rate reductions during cholinergic neuron photoactivation were consistent with graded reductions in heart rate with increased strength of electrical stimulation of the vagus nerve (Ng et al., 2001). Furthermore, our findings are consistent with the chronotropic cholinergic response observed during exogenous administration of ACh (Takahashi et al., 2003). ACh has been shown to induce a dose-dependent reduction of sinus rate of approximately 25-67% (Glukhov et al., 2010; Lang et al., 2011) in isolated mouse hearts, similar to our results of 46% maximal reduction. However, AV delay was considerably longer during such ubiquitous muscarinic activation (>24% (Lang et al., 2011)) compared to our approach, where we observed only up to 12% increase in AV conduction delay in some hearts. Our finding of no detectable change in P wave duration during photoactivation is also consistent with previous reports that found no change in right atrial conduction velocity during ACh perfusion (Schuessler et al., 1990).

Photoactivation using the micro LED, which provided the highest irradiance in our studies, produced a response similar to ubiquitous pharmacologic activation of muscarinic receptors, as observed with bethanechol. In addition to the effects of increased irradiance, the micro LED was more localized to the right atrial epicardium, having a high concentration of parasympathetic innervation, particularly at the RA/SVC junction (Ripplinger et al., 2016). Light from the micro LED could be oriented to selectively illuminate areas of the atria, such as the SA node, resulting in heart rate reductions without significant increases in AV delay, similar to the results of previous studies (Randall et al., 1986; Sampaio et al., 2003). On the other hand, non-selective illumination of the atria, that included the SA and AV nodes, by the spotlight LED resulted in a reduction of heart rate that was accompanied by an AV conduction delay.

Irradiances for optogenetic activation of cardiac tissues typically range from 0.5 to 10 mW/mm^2^, depending upon the cell type (Ambrosi et al., 2015; Bruegmann et al., 2010; Entcheva, 2013; Nyns et al., 2016). This range is consistent with the results of our LED spotlight and micro LED illumination. In future studies, the use of LED spotlight illumination could be helpful to confirm ChR2&EYFP expression, to generate quick HR responses. The micro LED lends itself to detailed study of the neuroanatomy of vagal inputs due to its more localized illumination. In addition, future devices for animals might soon be available to implant such micro LEDs into transgenic mice for *in-vivo* optogenetic activation of specific nerve populations (Kim et al., 2010; Xu et al., 2014).

### 4.1 Limitations

Our studies sought to examine intrinsic parasympathetic pathways and downstream responses in mouse hearts using an optogenetic approach to selective cholinergic neuron activation. We did not address specific cellular mechanisms at the post-synaptic level that control downstream signaling. Our studies focused on illumination of the RA/SVC junction – the presumed location of the sinoatrial node and did not focus in detail on the differences in the effect of left versus right sided parasympathetic stimulation. While directed at this general area, the focus of our spotlight LED was inexact. We believe this to be responsible for the variation in response demonstrated by the various irradiances studied using the spotlight LED and also for the induction of AV block, which could be a consequence of asynchronous or hyperparasympathetic stimulation, especially around the AV node (Ardell and Randall, 1986; Schiereck et al., 2000). Using a more targeted illumination approach, in future studies we plan to further elucidate the exact neuronal pathways responsible for parasympathetic responses. *In-vivo* studies may also be valuable to understand the whole-animal response to light activation of cardiac parasympathetic pathways.

## 5. Summary

Cardiac autonomic tone is a prevailing modulator of cardiac output and understanding the mechanisms of its control has taken center stage as the aging population suffers from degenerative cardiac disease. Existing electrical and pharmacological approaches to autonomic modulation, while effective in activating sympathetic and parasympathetic pathways, are non-selective proximal modulators. Optogenetic actuation of cardiac neurons provides a targeted, binary input with minimal cross-reactivity on distal cardiac tissue and is a powerful approach for in-depth examination of the endogenous interplay between the sympathetic and parasympathetic arms of the cardiac autonomic system.

We crossbred mice to express ChR2 in peripheral cholinergic neurons using Cre-Lox recombination driven by a choline acetyltransferase (ChAT) promoter. Upon illumination, excised perfused hearts from these animals exhibited sudden and dramatic reductions in heart rate with restoration of the previous heart rate after cessation of illumination. Such photoactivation of intrinsic cholinergic neurons reduced heart rate in an immediate, dose-dependent fashion, resembling the slowing of sinus rate in response to ACh. Our results demonstrate a new approach for controlling cardiac parasympathetic activity by selectively activating the endogenous release of ACh from intrinsic cardiac cholinergic neurons. This approach also enables spatiotemporal anatomic-functional probing of the intrinsic cardiac nervous system to reveal new insights that could be difficult to obtain using conventional approaches.

## Author contributions

AM, KE, MS, provided equal first-author contributions, designed and conducted the experiments, analyzed the data, and wrote the manuscript. AM, MS generated the final figures. MKD, KE, AM conducted the confocal imaging. MKD conducted immunohistochemistry and edited the manuscript. JB, KE conducted the optical mapping experiments. IRE, GT provided overall intellectual input and edited the manuscript. MWK, DM conceived the experiments, analyzed the data, provided overall intellectual input, and edited the manuscript.

## Acknowledgements

The microscopy expertise of Anastas Popratiloff, MD, PhD and the facilities GWU SEH Microscopy Core Facility are gratefully acknowledged. Matthew Stoyek, PhD is gratefully acknowledged for guidance in antibody staining of ChAT. Matthew Colonnese, PhD and Marnie Phillips, PhD are gratefully acknowledged for providing mice used in early experiments. We also thank Emilia Entcheva, PhD for many insightful discussions regarding photoactivation of channelrhodopsin within cardiac tissues.

